# The Northern Arizona SNP Pipeline (NASP): accurate, flexible, and rapid identification of SNPs in WGS datasets

**DOI:** 10.1101/037267

**Authors:** Jason W. Sahl, Darrin Lemmer, Jason Travis, James M. Schupp, John D. Gillece, Maliha Aziz, Elizabeth M. Driebe, Kevin Drees, Nathan Hicks, Charles H.D. Williamson, Crystal Hepp, David Smith, Chandler Roe, David M. Engelthaler, David M. Wagner, Paul Keim

## Abstract

Whole genome sequencing (WGS) of bacteria is becoming standard practice in many laboratories. Applications for WGS analysis include phylogeography and molecular epidemiology, using single nucleotide polymorphisms (SNPs) as the unit of evolution. The Northern Arizona SNP Pipeline (NASP) was developed as a reproducible pipeline that scales well with the large amount of WGS data typically used in comparative genomics applications. In this study, we demonstrate how NASP compares to other tools in the analysis of two real bacterial genomics datasets and one simulated dataset. Our results demonstrate that NASP produces comparable, and often better, results to other pipelines, but is much more flexible in terms of data input types, job management systems, diversity of supported tools, and output formats. We also demonstrate differences in results based on the choice of the reference genome and choice of inferring phylogenies from concatenated SNPs or alignments including monomorphic positions. NASP represents a source-available, version-controlled, unit-tested method and can be obtained from tgennorth.github.io/NASP.

## Introduction

Whole genome sequence (WGS) data from bacteria are rapidly increasing in public databases and have been used for outbreak investigations [1, 2], associating phylogeny with serology [3], as well as phylogeography [4]. WGS data are frequently used for variant identification, especially with regards to single nucleotide polymorphisms (SNPs). SNPs are used because they provide stable markers of evolutionary change between genomes [5]. Accurate and reliable SNP identification requires the implementation of methods to call, filter, and merge SNPs with tools that are version controlled, unit tested, and validated [6].

Multiple pipelines are currently available for the identification of SNPs from diverse WGS datasets, although the types of supported input files differ substantially. There are few pipelines that support the analysis of both raw sequence reads as well as genome assemblies. The ISG pipeline [7] calls SNPs from both raw reads, primarily from the Illumina platform, and genome assemblies, but wasn’t optimized for job management systems and only exports polymorphic positions. While only polymorphic positions may be adequate for many studies, including monomorphic positions in the alignment has been shown to be important for various phylogenetic methods. A commonly used SNP analysis software method is kSNP, which has been discussed in three separate publications [8-10]. The most recent version of kSNP (v3) doesn’t directly support the use of raw reads in the identification of SNPs. kSNP is a reference-independent approach in which all kmers of a defined length are compared to identify SNPs. The all-versus-all nature of the algorithm can result in a large RAM footprint and can stall on hundreds of bacterial genomes [7]. Finally, REALPHY was published as a method to identify SNPs using multiple references and then merging the results [11]. The authors claim that single reference based methods bias the results, especially from mapping raw reads against a divergent reference genome.

Additional methods have also been published that only support specific input formats. Parsnp is a method that can rapidly identify SNPs from the core genome, but currently only processes closely related genome assemblies [12]. SPANDx is a method that only supports raw reads, but does run on a variety of job management systems [13]. The program lyve-SET has been used in outbreak investigations and uses raw or simulated reads to identify SNPs [14]. Finally, the CFSAN SNP-pipeline is a published pipeline from the United States Food and Drug Administration that only supports the use of raw reads [15]. There have been no published comparative studies to compare the functionality of these pipelines on a range of test datasets.

In this study, we describe the Northern Arizona SNP Pipeline (NASP). NASP is a source-available, unit-tested, version-controlled method to rapidly identify SNPs that works on a range of job management systems, incorporates multiple read aligners and SNP callers, works on both raw reads and genomes assemblies, calls both monomorphic and polymorphic positions, and has been validated on a range of diverse datasets. We compare NASP with other methods, both reference-dependent and reference-independent, in the analysis of three reference datasets.

## Methods

NASP is implemented in Python and Go. NASP accepts multiple file formats as input, including “.fasta”, “.sam”, “.bam”, “.vcf”, “.fastq”, and “fastq.gz”. NASP can either function through a question/answer command line interface designed for ease of use, or through an argument-driven command-line interface. NASP was developed to work on job management systems including Torque, Slurm, and Sun/Oracle Grid Engine (SGE); a single node solution is available for NASP as well, but is not optimal. If filtering of duplicate regions in the reference genome is requested, the reference is aligned against itself with NUCmer [16]. These duplicated regions are then masked from downstream analyses, although are still available for investigation. If external genome assemblies are supplied, they are also aligned against the reference genome with NUCmer and SNPs are identified by a direct one-to-one mapping of the query to the reference. In the case of duplications in the query but not the reference, all copies are aligned and any differences at any given base are masked with an “N” character to identify it as ambiguous.

If raw reads are supplied, they can be adapter and/or quality trimmed with Trimmomatic [17]. Raw or trimmed reads are aligned against a FASTA-formatted reference using one of the supported short read aligners, including BWA-MEM [18], Novoalign, bowtie2 [19] and SNAP [20]. A binary alignment map (BAM) file is created with Samtools [21] and SNPs can be called with multiple SNP callers, including the UnifiedGenotyper method in GATK [22, 23], Samtools, SolSNP (http://sourceforge.net/projects/solsnp/), and VarScan [24]. Positions that fail a user-defined depth and proportion threshold are filtered from downstream analyses but are retained in the “master” matrices. A workflow of the NASP pipeline is shown in Figure 1 and a summary is shown in Supplemental Table 1.

**Figure 1.**
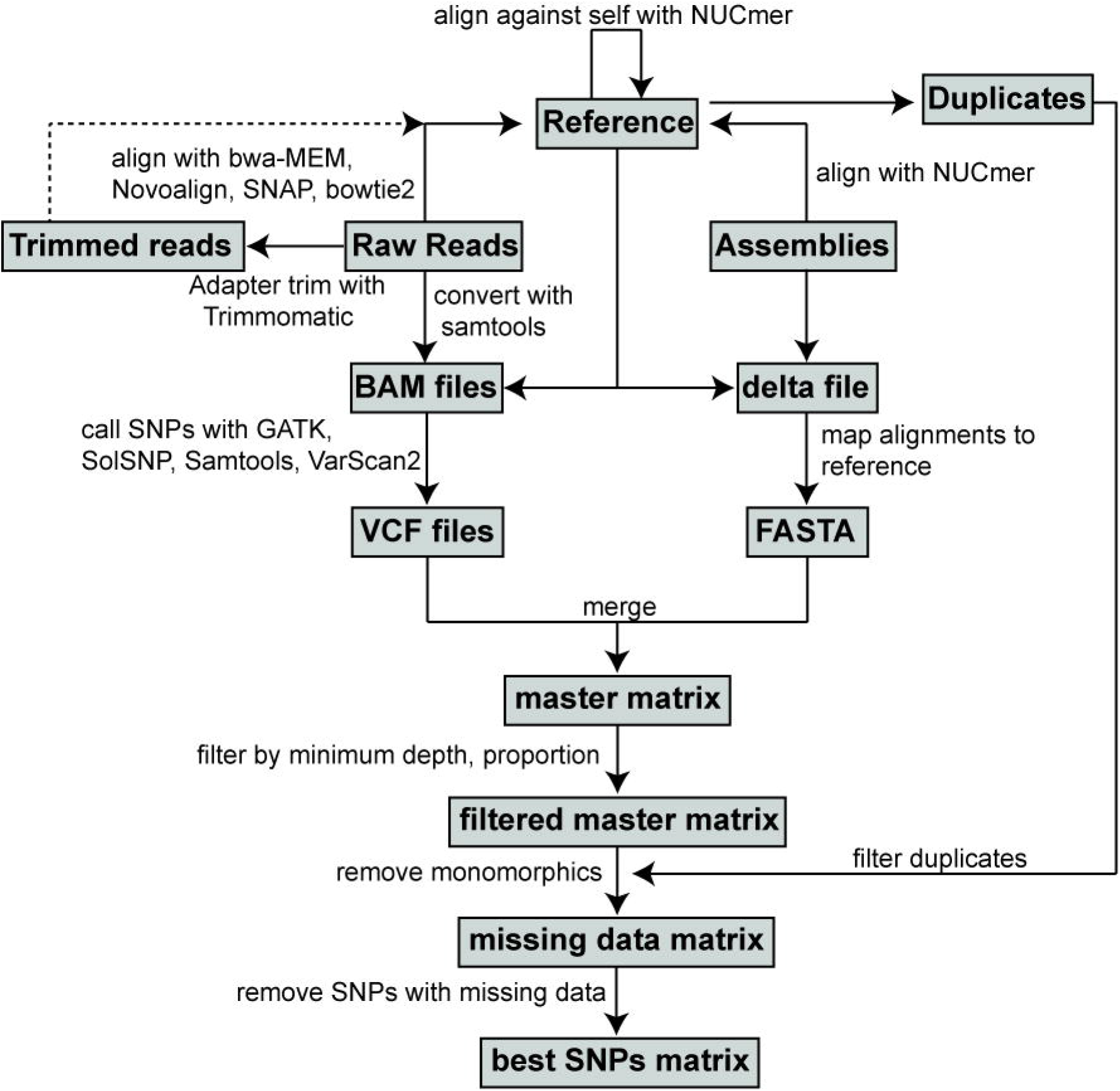
A workflow of the NASP algorithm. Optional steps are shown by dashed lines.

**Table 1.** An overview of commonly used SNP pipelines

| Pipeline name | Supported data types | Output type | Parallel Job management support? |
| --- | --- | --- | --- |
| NASP | FASTA,BAM,SAM,VCF,FASTQ,FASTQ.GZ | matrix, VCF, FASTA | SGE,SLURM,TORQUE |
| ISG | FASTA,BAM,VCF,FASTQ,FASTQ.GZ | matrix, FASTA | No |
| Parsnp | FASTA | gingr file, phylogeny, FASTA, VCF | No |
| REALPHY | FASTA*, FASTQ, FASTQ.GZ | multi-FASTA, phylogeny | No |
| SPANDx | FASTQ.GZ | Nexus file | SGE,SLURM,TORQUE |
| CFSAN | FASTQ, FASTQ.GZ | SNP list, FASTA | SGE,TORQUE |
| kSNPv3 | FASTA | Matrix, FASTA, phylogeny | No |
| Mugsy | FASTA | MAF file | No |
| LYVE-set | FASTQ.GZ, FASTA* | matrix, FASTA, phylogeny | SGE |

\*generates simulated reads

The results of the pipeline can include up to four separate SNP matrices. The first matrix is the master matrix (master.tsv), which includes all calls, both monomorphic and polymorphic, across all positions in the reference with no positions filtered or masked; positions that fall within duplicated regions are shown in this matrix, although they are flagged as duplicated. An optional second matrix (master_masked.tsv) can also be produced. This matrix is the same as the master matrix, although any position that fails a given filter (minimum depth, minimum proportion) is masked with an “N”, whereas calls that could not be made are given an “X”; this matrix could be useful for applications where all high-quality, un-ambiguous positions should be considered. The third matrix (missingdata.tsv) includes only positions that are polymorphic across the sample set, but can include positions that are missing in a subset of genomes and not found in duplicated regions; these SNPs have also been processed with the minimum depth and proportion filters and are still high quality calls. The last matrix (bestsnp.tsv) is a matrix with only polymorphic, non-duplicated, clean calls (A,T,C,G) that pass all filters across all genomes. FASTA files are automatically produced that correspond to the bestsnp and missingdata matrices.

In addition to the matrices and FASTA files, NASP produces statistics that can be useful for the identification of potentially problematic genomes, such as low coverage or mixtures of multiple strains. These statistics can also be used for determining the size of the core genome, including both monomorphic and polymorphic positions, of a given set of genomes.

Post matrix scripts are included with NASP in order to convert between file formats, remove genomes and/or SNPs, provide functional SNP information, and to convert into formats that can be directly accepted by other tools, such as Plink [25], a method to conduct genome wide association studies (GWAS). Documentation for all scripts is included in the software repository.

**Test datasets**. To demonstrate the speed and functionality of the NASP pipeline, three datasets were selected. The first includes a set of 21 *Escherichia coli* genome assemblies used in other comparative studies [11, 26] (Supplemental Table 2). REALPHY was run on self-generated single-ended simulated reads, 100bp in length. Additional pipelines were run with paired-end reads generated by ART chocolate cherry cake [27], using the following parameters: -l 100 -f 20 -p -ss HS25 -m 300 -s 50. Unless otherwise noted, the reference genome for SNP comparisons was K-12 MG1655 (NC_000913) [28]. All computations were performed on a single node, 16-core server with 48Gb of available RAM. For kSNP, the optimum k value was selected by the KChooser script included with the repository.

**Table 2.** SNP calling results on a set of 21 *E. coli* genomes

| Method | Reference | Data Type | Parameters | #SNPs considered | #Total sites | Walltime (single node - 8 cores) | Topological score | #defining SNPs |
| --- | --- | --- | --- | --- | --- | --- | --- | --- |
| NASP | K12 MG1655 | Assemblies | Default | 267978<sup>a</sup> | 2322434 | 10m00s | 100% | 809 |
| NASP | K12 MG1655 | Assemblies | NUCmer (-b 20) | 162758<sup>a</sup> | 1839583 | 10m00s | 100% | 744 |
| NASP | K12 MG1655 | ART PE reads | BWA, GATK, MinDepth=3, MinAF=0.90 | 170208<sup>a</sup> | 1984510 | 1h43m00s | 100% | 826 |
| NASP | *E. fergusonii* 35469 | Assemblies | Default | 244262<sup>a</sup> | 2227038 | 10m00s | 100% | 741 |
| NASP | *E. fergusonii* 35469 | ART PE reads | BWA, GATK, MinDepth=3, MinAF=0.90 | 141238<sup>a</sup> | 1813349 | 1h17m10s | 100% | 748 |
| ISG | K12 MG1655 | Assemblies | Default | 268524<sup>a</sup> | N/A | 6m47s | 100% | 810 |
| ISG | K12 MG1655 | ART PE reads | minaf 0.9, mindp 3 | 206193<sup>a</sup> | N/A | 14m45s | 100% | 824 |
| Parsnp | K12 MG1655 | Assemblies | "-c d" | 151256<sup>a</sup> | 1682404 | 4m35s | 100% | 777 |
| REALPHY | K12 MG1655 | REALPHY SE reads | Default | 171828<sup>a</sup> | 1897146 | 3h11m00s | 100% | 779 |
| kSNPv3 | N/A | Assemblies | -core | 20587<sup>a</sup> | N/A | 27m58s | 91.80% | 5 |
| kSNPv3 | N/A | Assemblies | Default | 284134 | N/A | 27m58s | 95.80% | 547 |
| Spandx | K12 MG1655 | ART PE reads | Default | 95214<sup>a</sup> | N/A | N/A | 100% | 709 |
| CFSAN | K12 MG1655 | ART PE reads | Default | 128512<sup>a</sup> | N/A | 1h56m00s | 100% | 808 |
| Mugsy | N/A | Assemblies | Default | 307072<sup>a</sup> | 2478794 | 1h39m03s | 100% | unknown |
| lyve-SET | K12 MG1655 | ART PE reads | min_coverage 3, min_alt_frac 0.9 | 163118<sup>a</sup> | 1183153 | 6h25m | 85% | 329 |

<sup>a</sup>strictly core genome SNPs

The second dataset includes a set of 15 *Yersinia pestis* genomes from North America (Supplemental Table 3). For those external SNP pipelines that only support raw reads, simulated reads were generated from genome assemblies with ART. A set of SNPs (Supplemental Table 4) has previously been characterized on these genomes with wet-bench methods (unpublished). This set was chosen to determine how many verified SNPs could be identified by different SNP pipelines. All computations were performed on a single node, 16-core server with 48Gb of available RAM.

**Table 3.** SNP calling results on a set of *Y. pestis* genomes

| Method | Data type | Parameters | #called SNPs | #CO92 errors (n=13) | #verified SNPs (n=26) | Vital SNF (n=9) |
| --- | --- | --- | --- | --- | --- | --- |
| NASP | ART simulated reads | BWA, GATK, MinDepth=3, MinAF=0.90 | 147 | 13 | 26 | 9 |
| NASP | assemblies | default | 181 | 13 | 26 | 9 |
| ISG | ART simulated reads | minaf=3, mindp = 0.9 | 151 | 13 | 26 | 9 |
| ISG | assemblies | default | 177 | 13 | 26 | 9 |
| Parsnp | assemblies | default | 141 | 12 | 23 | 7 |
| REALPHY | REALPHY simulated reads | default | 163 | 12 | 25 | 9 |
| SPANDx | ART simulated reads | default | 150 | 13 | 25 | 9 |
| kSNPv3 | assemblies | k=19 | 130 | 11 | 21 | 5 |
| CFSAN | ART simulated reads | default | 250 | 13 | 26 | 9 |
| lyve-SET | ART simulated reads | min_coverage 3, min_alt_frac 0.9 | 402 | 13 | 26 | 9 |

**Table 4.** Simulated data results

| Method | Data type | #called SNPs | SNPs in duplicated regions | Filtered SNPs | Total SNPs | Topologica score |
| --- | --- | --- | --- | --- | --- | --- |
| NASP | simulated reads | 3202 | 232 | 67 | 3501 | 98.50% |
| NASP | simulated assemblies | 3269 | 232 | N/A | 3501 | 98.50% |
| Parsnp | simulated assemblies | 3492 | unknown | N/A | 3492 | 95.60% |
| ISG | simulated reads | 3258 | 126 | 8 | 3392 | 92.40% |
| ISG | simulated assemblies | 3266 | 235 | N/A | 3501 | 95.60% |
| SPANDx | simulated reads | 2132 | unknown | 116 | 2248 | 87.20% |
| CFSAN | simulated reads | 3290 | unknown | unknown | 3290 | 95.30% |
| REALPHY | simulated assemblies | 3320 | unknown | unknown | 3320 | 91.60% |
| kSNPv3 | simulated assemblies | 3304 | unknown | N/A | 3304 | 91.90% |
| lyve-SET | simulated reads | 3460 | unknown | unknown | 3460 | 95.80% |

The last dataset includes simulated data from *Yersinia pestis.* Reads and assemblies from 133 *Y. pestis* genomes [29] were downloaded from public databases and processed with NASP using the CO92 genome as the reference to produce a reference phylogeny for WGS data simulation. Assemblies and reads were simulated from this reference phylogeny and a reference genome (CO92 chromosome) using TreeToReads (https://github.com/snacktavish/TreeToReads), introducing 3501 mutations. A phylogeny was inferred from the concatenated SNP alignment (3501 simulated SNPs produced by TreeToReads) with RAxML v8 to provide a ‘true’ phylogeny for the simulated data. Simulated reads (250bp) and assemblies were both processed with pipelines to identify how many of these introduced SNPs could be identified.

To test the scalability of NASP on genome assemblies, a set of 3520 *E. coli* genomes was selected (Supplemental Table 5). Genomes were randomly selected with a python script (https://gist.github.com/jasonsahl/990d2c56c23bb5c2909d) at various levels (100-1000) and processed with NASP. In this case, NASP was run on multiple nodes across a 31-node cluster at Northern Arizona University. The elapsed time was reported only for the step where aligned files are compiled into the resulting matrix. Time required for the other processes is dependent on the input file type and the amount of available resources on a HPC cluster.

**External SNP pipelines**. Multiple SNP pipelines, both reference-dependent and reference-independent, were compared with NASP, including kSNP v3.9.1 [10], ISG v0.16.10-3 [7], Parsnp v1.2 [12], REALPHY v112 [11], SPANDx v2.7 [13], Mugsy v1r2.2 [30], lyve-SET v1.1.6 [31], and CFSAN (https://github.com/CFSAN-Biostatistics/snp-pipeline). Exact commands used to run each method are shown in Supplemental Data File 1. An overview of all tested methods is shown in Table 1. Most of the methods output FASTA files, which were used to infer phylogenies. For Mugsy, the MAF file was converted to FASTA with methods described previously [32].

**Phylogenetics**. Phylogenies were inferred using a maximum likelihood algorithm implemented in RAxML v8.1.7 [33], except where noted. The exact commands used to infer the phylogenies are shown in Supplemental Data File 1. Tree topologies were also compared on the same input data. Commands to infer these phylogenies using FastTree2 [34], ExaBayes [35], and Parsimonator (github.com/stamatak/Parsimonator-1.0.2) are shown in Supplemental Data File 1.

**Dendrogram of multiple methods**. To visually represent how well different methods relate, a dendrogram was generated. Each phylogeny was compared against a maximum likelihood phylogeny inferred from the reference test set with compare2trees. A UPGMA dendrogram was then calculated with Phylip [36] on the resulting similarity matrix.

## Results

**Pipeline functionality and post-matrix scripts**. NASP is a reference-dependent pipeline that can incorporate both raw reads and assemblies in the SNP discovery process; NASP was not developed for the identification and annotation of short insertions/deletions (indels). NASP can use multiple aligners and SNP callers to identify SNPs and the consensus calls can be calculated across all methods. An additional strength of NASP is that it can run on multiple job management systems as well as on a single node. A complete workflow of the NASP method is shown in Figure 1. Several post-matrix scripts are included with NASP in order to convert between file formats, including generating input files for downstream pipelines, including Plink [25]. An additional script can annotate a NASP SNP matrix using SnpEff [37] to provide functional information for each SNP.

**NASP run time scalability**. To visualize how NASP scales on processing genome assemblies, a set of 3520 *E. coli* genomes was sampled at 100 genome intervals and processed with NASP with 10 replicates. The results demonstrate that the matrix building step in NASP scales linearly with the processing of additional genomes (Figure 2A). The memory footprint of this step also scales linearly (Figure 2B) and doesn’t exceed 4Gb on a large set of genomes (n=1000). If raw reads are used, additional time is required for the alignment and SNP calling methods, and the overall wall time would scale with the number of reads that needed to be processed. The matrix-building step, where assemblies and VCF files are merged into the matrix, would scale linearly regardless of the SNP identification method chosen.

**Figure 2.**
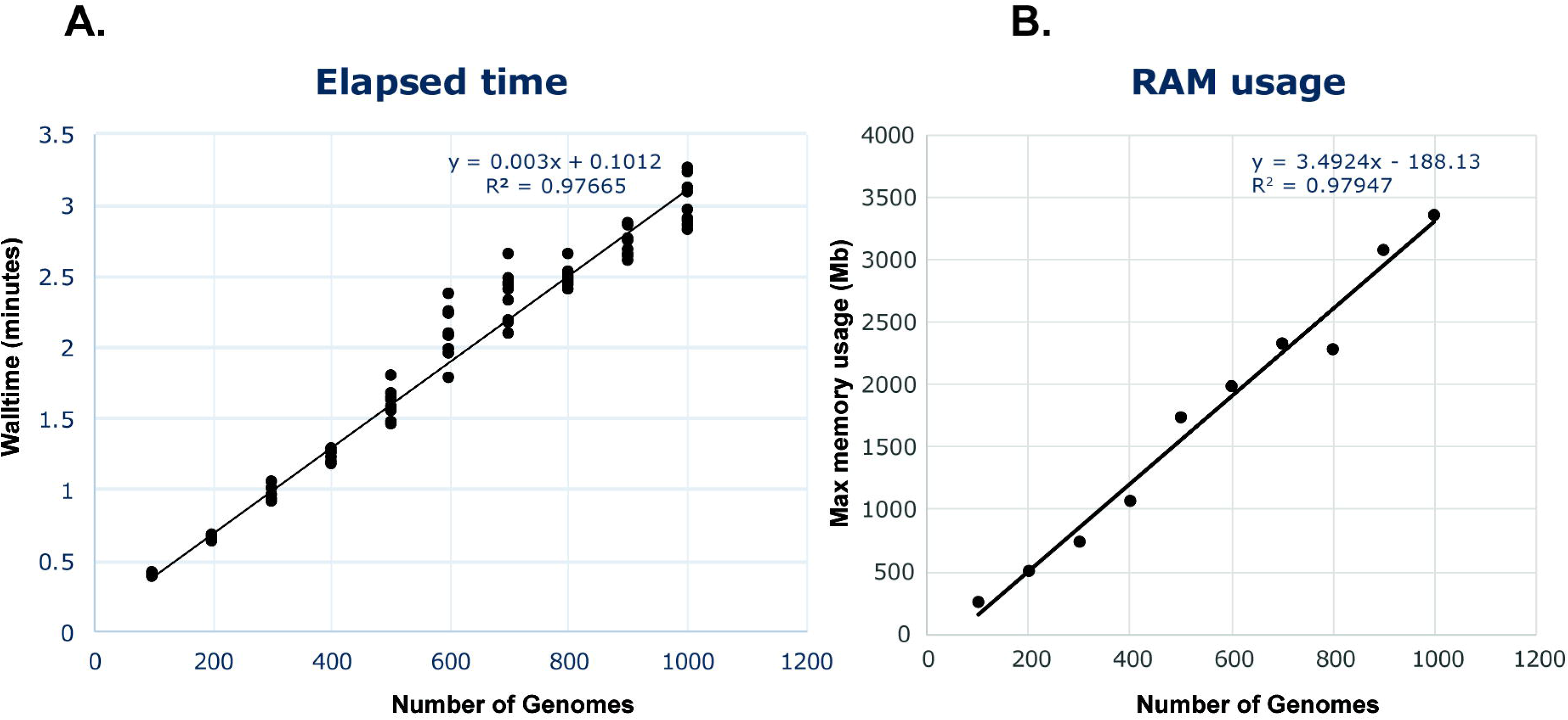
NASP benchmark comparisons of walltime (A) and RAM (B) on a set of *Escherichia coli* genomes. For the walltime comparisons, 3520 *E. coli* genomes were randomly sampled ten times at different depths and run on a server with 856 cores. Only the matrix building step is shown, but demonstrates a linear scaling with the processing of additional genomes.

**Pipeline comparisons on *E. coli* genomes data set**. To test differences between multiple pipelines, a set of 21 *E. coli* genomes used in other comparative genomics studies [11, 26] were downloaded and processed with Parsnp, SPANDx, kSNPv3, ISG, REALPHY, CFSAN, lyve-SET, Mugsy, and NASP. For methods that do not support genome assemblies, paired end reads were simulated with ART, while single end reads were used by REALPHY, as this method is integrated into the pipeline.

To identify how well the simulated paired end reads represent the finished genomes, a NASP run was conducted on a combination of completed genome assemblies as well as simulated raw reads. The phylogeny demonstrates that assemblies and raw reads fall into identical locations (Supplemental Figure 1), suggesting that the paired end reads are representative of the finished genome assemblies.

The authors of REALPHY assert that their analysis of this dataset demonstrates the utility of using their approach to avoid biases in the use of a single reference genome by using multiple references [11]. However, in our tests, we could only get REALPHY to complete when using a single reference. To test differences between methods, SNPs were identified with multiple reference-dependent and -independent methods, and maximum likelihood (ML) phylogenies were compared. The results demonstrate that all methods, with the exception of kSNPv3 and lyve-SET, returned a phylogeny with the same topology as the published phylogeny [11] (compare2trees topological score = 100%) (Table 2). The run wall time demonstrates that most other methods were significantly faster than REALPHY (Table 2), even when REALPHY was called against a single reference. Wall time comparisons between methods are somewhat problematic, as some pipelines infer phylogenies and others, including NASP, do not. Additionally, using raw reads is generally expected to be slower than using a draft or finished genome assembly. Finally, some methods are optimized for job management systems, whereas others were designed to run on a single node. For these comparisons, all methods that have single node support were run on a single node. Only SPANDx seems to be dependent on job management systems and could not be successfully run on a single node.

One of the other assertions of the REALPHY authors is that phylogenies reconstructed using an alignment of concatenated SNPs are unreliable [11, 26], especially with regards to branch length biases [38]. However, the phylogeny inferred from a NASP alignment of monomorphic and polymorphic sites was in complete agreement with the topology of the phylogeny inferred from a concatenation of only SNPs (compare2trees topological score = 100%); tree lengths were indeed variable by using these two different input types using the same substitution model (Supplemental Figure 2). We also employed an ascertainment bias correction (Lewis correction) implemented in RaxML [38], in order to correct for the use of only polymorphic sites, and found no difference between tree topologies using substitution models that did not employ this correction (data not shown). For this dataset of genomic *E. coli* assemblies, there appears to be no effect of using a concatenation of polymorphic sites on the resulting tree topology, although branch lengths were affected compared to an alignment containing monomorphic sites.

To understand how the choice of the reference affects the analysis, NASP was also run using *E. coli* genome assemblies and simulated reads against the outgroup, *E. fergusonii,* as the reference. The results demonstrate that the same tree topology was obtained by using a different, and much more divergent, reference (compare2trees topology score = 100%). However, in both cases, fewer SNPs were identified by using a divergent reference (Table 2).

Some authors suggest that reference-independent approaches are less biased and more reliable than reference dependent-approaches [8]. For the case of this *E. coli* dataset, the phylogeny inferred by Mugsy, a reference-independent approach, was in topological agreement with other reference-dependent approaches (Table 2). In fact, kSNPv3 was one of the only methods that returned a tree phylogeny that was inconsistent with all other methods (Table 2); an inconsistent kSNP phylogeny has also been reported in the analysis of other datasets [15]. To analyze this further, we identified SNPs (n=826) from the NASP run using simulated paired-end reads that were uniquely shared on a branch of the phylogeny that defines a monophyletic lineage (Supplemental Figure 3). We then calculated how many of these SNPs were identified by all methods and found widely variable results (Table 2). Using kSNP with only core genome SNPs identified only 5 of these SNPs, which explains the differences in tree topologies.

**Figure 3.**
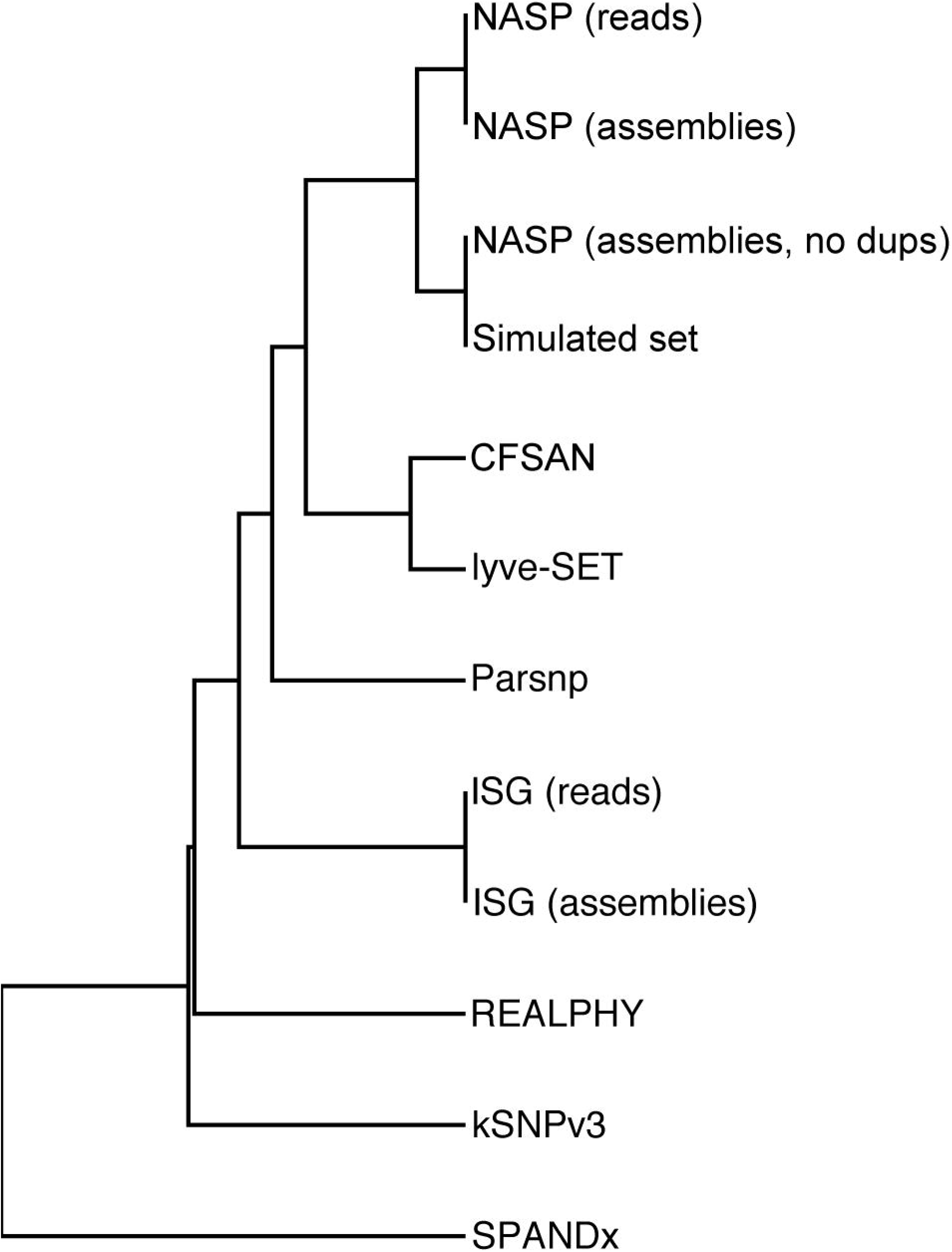
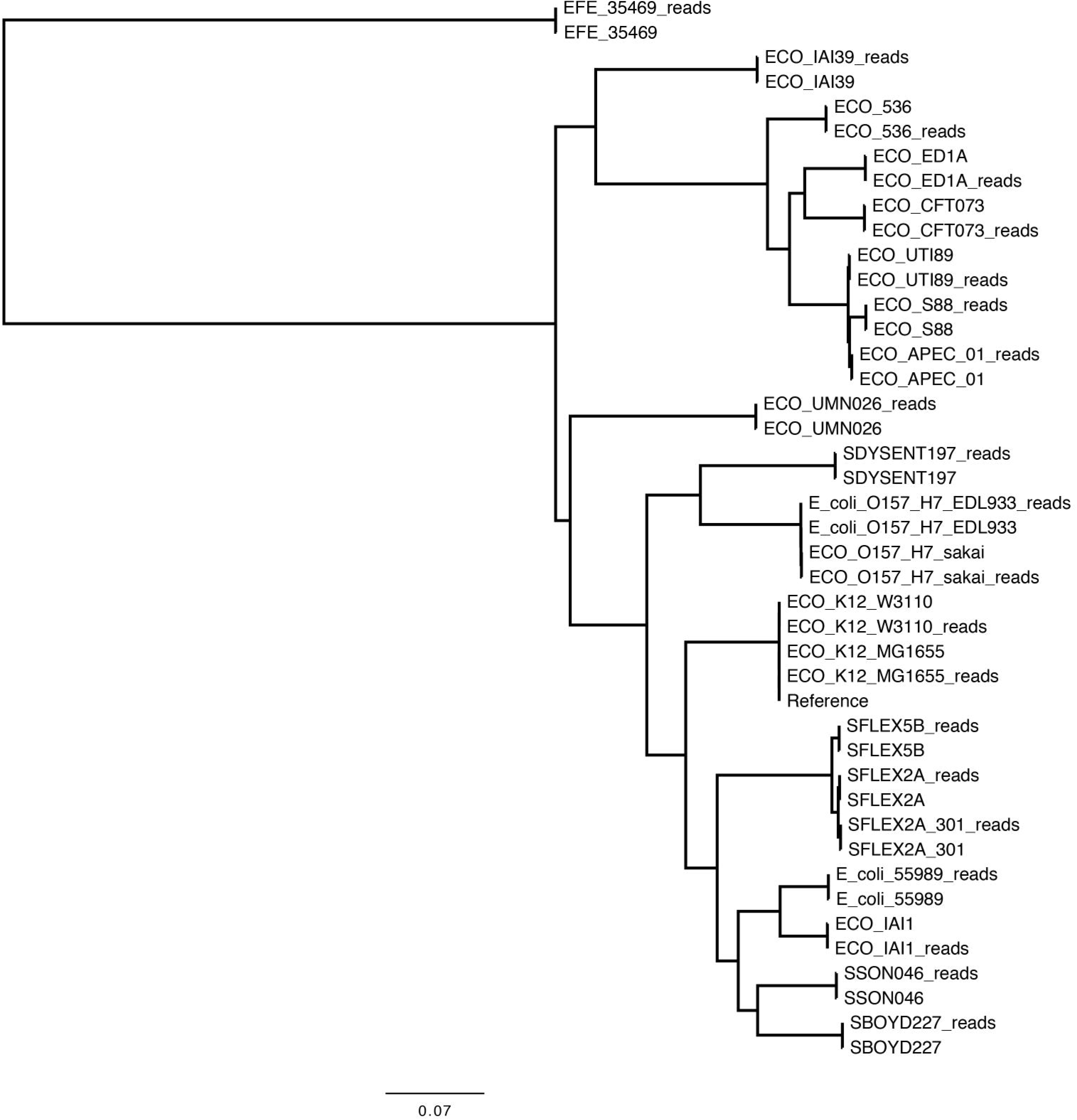
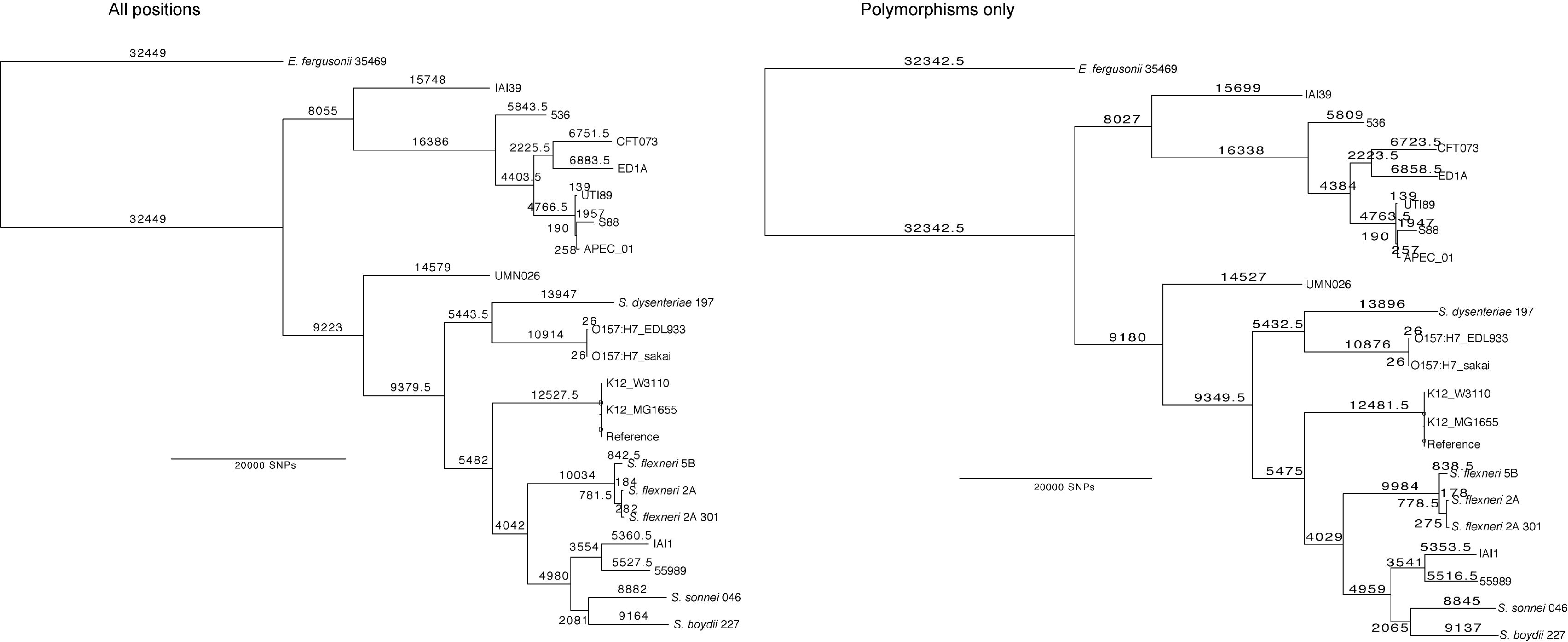
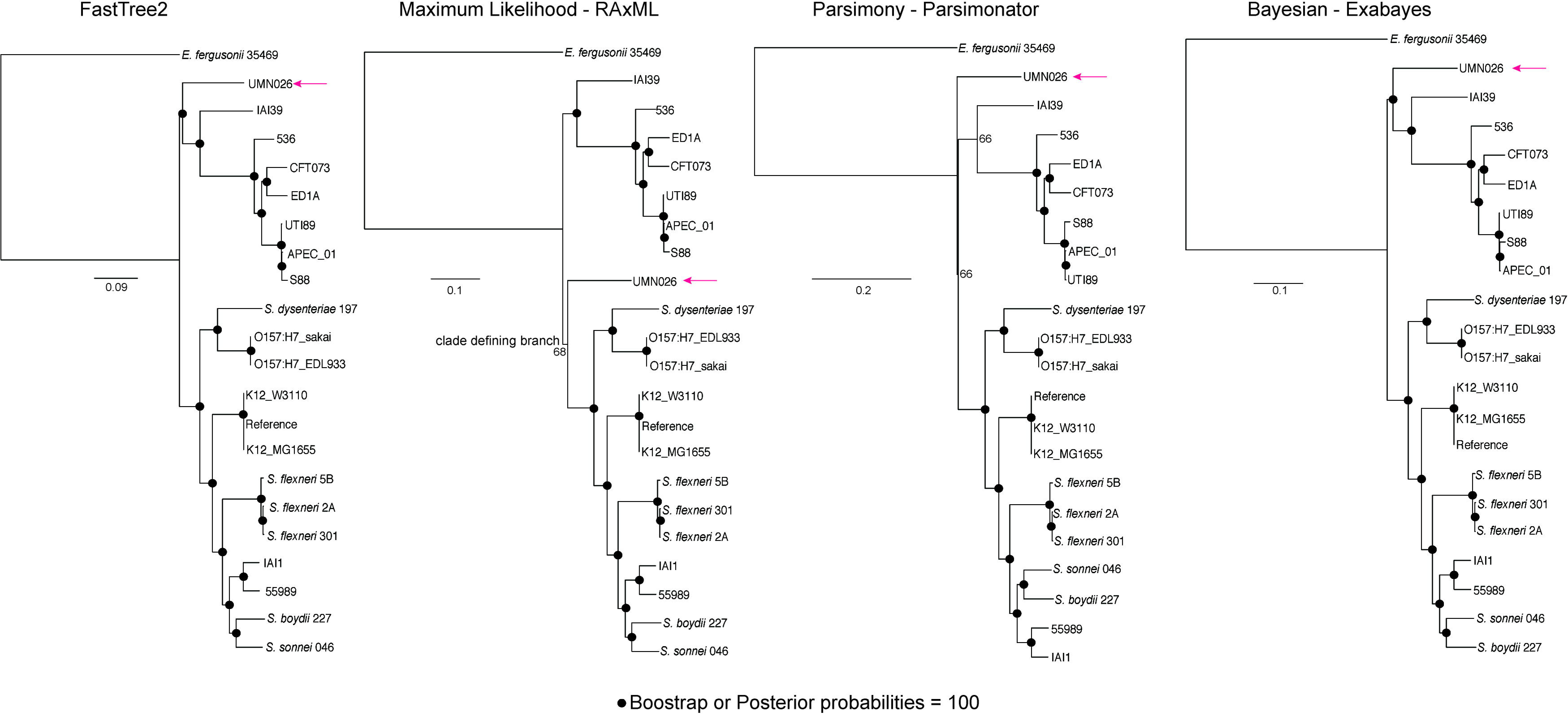
Dendrogram of tree building methods on a simulated set of mutations in the genome of *Yersinia pestis* Colorado 92. The topological score was generated by compare2trees compared to a maximum likelihood phylogeny inferred from a set of 3501 SNPs inserted by Tree2Reads. The dendrogram was generated with the Neighbor method in the Phylip software package [36].

In many cases, the same tree topology was returned even though the number of identified SNPs differed dramatically (Table 2). This result could be due to multiple factors, including if and how duplicates are filtered from the reference genome or other genome assemblies. With regards to NASP, erroneous SNPs called in genome assemblies are likely artifacts from the whole genome alignments using NUCmer. The default value for aligning through poorly scoring regions before breaking an alignment in NUCmer is 200, potentially introducing many spurious SNPs into the alignment, especially in misassembled regions in draft genome assemblies. By changing this value to 20, the same tree topology was obtained, although many fewer SNPs (n=~100,000) were identified (Table 2). This value is easily altered in NASP and should be tuned based on the inherent expected diversity in the chosen dataset. Additional investigation is required to verify that SNPs in divergent regions are not being lost by changing this parameter. Another option is to use simulated reads from the genome assemblies in the SNP identification process.

**Phylogeny differences on the same dataset**. Previously, it has been demonstrated that different phylogenies can be obtained on the same dataset using either RAxML or FastTree2 [15]. To test this result across multiple phylogenetic inference methods, the NASP *E. coli* read dataset was used. Phylogenies were then inferred using a maximum likelihood method in RAxML, a maximum parsimony method implemented in Parsimonator, a minimum evolution method in FastTree2, and a Bayesian method implemented in Exabayes [35]. The results demonstrate variability in the placement of one genome (UMN026) depending on the method. FastTree2 and Exabayes agreed on their topologies, including 100% congruence of the replicate trees. The maximum likelihood and maximum parsimony phylogenies were slightly different (Supplemental Figure 3) and included low bootstrap replicate values at the variable node. The correct placement of UMN026 is unknown and is likely confounded by the extensive recombination observed in *E. coli* [39].

**Pipeline comparisons on a well characterized dataset**. To test the functionality of different SNP calling pipelines, a set of 15 finished *Yersinia pestis* genomes were compared. This set of genomes was selected because 26 SNPs in the dataset have been verified by wet-bench methods (Supplemental Table 4). Additionally, 13 known errors in the reference genome, *Y. pestis* CO92 [40], have been identified (Supplemental Table 4) and should consistently be identified in SNP discovery methods. The small number of SNPs in the dataset requires accurate SNP identification to resolve the phylogenetic relationships of these genomes.

The results demonstrate differences in the total number of SNPs called between different methods (Table 3). Most of the methods identified all 13 known sequencing errors in CO92, although Parsnp, REALPHY, and kSNPv3 failed to do so. The number of verified SNPs also varied between methods, from 21 in kSNPv3 to all 26 in multiple methods (Table 3). An analysis of wet-bench validated SNPs (n=9) that are identified in more than one genome demonstrated that some methods failed to identify all of these SNPs, which could lead to a very different phylogeny than the phylogeny using these SNPs that are vital for resolving important phylogenetic relationships. These SNPs could represent differences that could differentiate between strains in an outbreak event.

**Pipeline comparisons on a simulated set of assemblies and reads**. Simulated data for *Y. pestis* were used to compare SNP identification between pipelines. In this method, 3501 mutations (Supplemental Data File 2) were inserted into genomes based on a published phylogeny [41] and FASTA file. Raw reads were also simulated from these artificially mutated assemblies with ART to generate paired end sequences. Reads and assemblies were run across all pipelines, where applicable.

The results demonstrate that NASP identified all of the inserted SNPs using raw reads, although 68 SNPs failed the proportion filter (0.90) and 232 SNPs fell in duplicated regions (Table 4); some of the duplicated SNPs would also fail the proportion filter. Of all other methods, only ISG identified all inserted mutations. SPANDx only identified 2248 SNPs when run with default values. Parsnp identified the majority of the mutations, although duplicate regions appear to have also been aligned.

To understand how the SNPs called would affect the overall tree topology, a phylogeny was inferred for each set of SNPs with RAxML. A similarity matrix was made for each method based on the topological score compared to the ML phylogeny inferred from the known mutations. The UPGMA dendrogram demonstrates that the NASP results return a phylogeny that is more representative of the “true” phylogeny than other methods (Figure 3). Without removing SNPs found in duplicated regions, the NASP phylogeny was identical to the phylogeny inferred from the known SNPs.

## Discussion

Understanding relationships between bacterial isolates in a population is important for applications such as source tracking, outbreak investigations, phylogeography, population dynamics, and diagnostic development. With the large numbers of genomes that are typically associated with these investigations, methods are required to quickly and accurately identify SNPs in a reference population. However, no studies have conducted a broad analysis to compare published methods on real and simulated datasets to identify relevant strengths and weaknesses.

Multiple publications have used a reference-dependent approach to identify SNPs to understand population dynamics [38]. While the specific methods are often published, the pipelines to run these processes are often un-published [42, 43], which complicates the ability to replicate results. NASP has already been used to identify SNPs from multiple organisms, including fungal [44] and bacterial [45, 46] pathogens. The version-controlled source code is available for NASP, which should ensure the replication of results across research groups.

Recently it has been suggested that the use of a single reference can bias the identification of SNPs, especially in divergent references [11]. In our *E. coli* test set, ~29,000 fewer SNPs were called by aligning *E. coli* reads against the reference genome of the outgroup, *E. fergusonii,* compared to the K-12 reference, although the tree topologies were identical (Table 2). In the *E. coli* test set phylogeny, the major clades are delineated by enough SNPs that the loss of a small percentage is insufficient to change the overall tree topology, although the branch lengths were variable. In other datasets, the choice of the reference should be made carefully to include as many SNPs as needed to define the population structure of a given dataset.

According to the authors of kSNP, a k-mer-based reference-independent approach, there are times where alignments are not appropriate in understanding bacterial population structure [8]. In our *E. coli* analysis, reference-dependent and reference-independent methods generally returned the same tree topology (Table 2), with the exception of kSNPv3 and lyve-SET, using only core genome SNPs. Using all of the SNPs identified by kSNPv3 also gave a different tree topology than the other methods (Table 2). A detailed look at branch specific SNPs demonstrated that using kSNP with core SNPs failed to identify most of the branch specific SNPs for one of the major defining clades (Table 2). For datasets that are only defined by a small number of SNPs, a method should be chosen that includes as many SNPs as possible in order to maximize the relevant search space. While NASP cannot truly use the pan-genome if a single reference genome is chosen, it can incorporate data from all positions in the reference genome if missing data are included in the alignment. A true pan-genome reference can be used with NASP to more comprehensively identify SNPs, but curation of the pan-genome is necessary to remove genomic elements introduced by horizontal gene transfer that could potentially confound the phylogeny.

Phylogenetics on an alignment of concatenated SNPs is thought to be less preferable than an alignment that also contains monomorphic positions [11, 38]. However, the inclusion of monomorphic positions can drastically increase the run time needed to infer a phylogeny, especially where the population structure of a species can be determined by a small number of polymorphisms. Substitution models are available in RAxML v8 that contain acquisition bias corrections that should be considered when inferring phylogenies from concatenated SNP alignments. In our *E. coli* test case, using concatenated SNPs did not change the tree topology compared to a phylogeny inferred from all sites, but did affect branch lengths (Supplemental Figure 2). For downstream methods that depend on accurate branch lengths, decisions must be made on whether or not to include monomorphic positions into the alignment. NASP provides the user with the flexibility to make those decisions in a reproducible manner.

NASP represents a version-controlled, unit-tested pipeline for identifying SNPs from datasets with diverse input types. NASP is a high throughput method that can take a range of input formats, can accommodate multiple job management systems, can use multiple read aligners and SNP callers, can identify both monomorphic and polymorphic sites, and can generate core genome statistics across a population.

